# Beyond brain size: disentangling the effect of sex and brain size on brain morphometry and cognitive functioning

**DOI:** 10.1101/2024.06.20.599908

**Authors:** Aliza Brzezinski-Rittner, Roqaie Moqadam, Yasser Iturria-Medina, Mallar Chakravarty, Mahsa Dadar, Yashar Zeighami

## Abstract

It is imperative to study sex differences in brain morphology and function. However, there are major observable and unobservable confounding factors that can contribute to the estimated differences. Males have larger head sizes than females. Head size differences not only act as a confounding factor in studying sex differences in the brain, but also impact its anatomy and functioning. In this work, we seek to disentangle the effect of head size from sex in studying sex differentiated aging trajectories, its relation to canonical functional networks and cytoarchitectural classes, brain allometry, cognition. Using the UK Biobank (UKBB) neuroimaging data (N = 35,732 participants, 19,281 females, 44-82 years of age), we created a subsample (N = 11,294) where females (N = 5,657) and males were matched by their total intracranial volume (TIV) and age, a subsample that maintains the UKBB sample distribution, one matched only by age, and one that exaggerated the TIV difference between sexes. We then modeled the aging trajectories at both regional and vertex-wise levels in the four subsamples, and compared the estimations of the models. Our results show that when females and males have the same head size, the overall sex estimations tend towards zero, suggesting that most of the variability results from head size differences. Our approach also revealed bidirectional sex differences in brain neuroanatomy previously masked by the effect of head size. Further, the scaling relationship between regional and total brain volume remains fairly consistent across the lifespan and is not sex differentiated overall. We evaluated how the results of cognitive tests with perceived sex differences are influenced and explained by head size and found that although the correlation between TIV and cognitive scores is low, the matching process changes the direction of the effect sizes of differences between sexes in “verbal and numerical reasoning” and “working memory” cognitive domains. Taken together, employing a matching approach that is widely used in causal modeling studies, we provide new evidence for disentanglement of sex differences in the brain from head size as a biological confound.

## Introduction

Aging is a ubiquitous biological process that is experienced by all individuals and plays a role as a primary risk factor for several brain disorders, particularly dementia ^1^. From a biological standpoint, aging results in molecular and cellular alterations ^2^ that entail various behavioral manifestations including change in cognition. Numerous studies on brain aging have provided evidence that aging leads to loss of cortical and subcortical gray and white matter volumes ^3–5^, decrease in cortical thickness ^6–8^, changes in white matter tracts ^9^, and an increase in white matter hyperintensity burden.^10^

Sex as a biological factor affects the aging process, with males showing an accelerated epigenetic age compared to females ^11^. There are established morphological brain differences between sexes across the lifespan ^12^, both globally and at a regional level ^13^. For example, when a 2 standard deviation change in microstructural and volumetric MRI measures compared to age of 45 was defined as a marker of aging, males showed an accelerated brain aging in several domains (e.g., total brain and hippocampal volumes, mean diffusivity) ^14^. Furthermore, neurodegenerative disorders such as Alzheimer’s disease (AD) and Parkinson’s disease (PD) are sex differentiated and both present with sex-specific clinical manifestations. Prevalence of AD has a female:male prevalence ratio of 2:1 ^15^ whereas the reported male:female incidence ratios of PD in Western and South American Populations ranges from 1.3 to 2.0 ^16^.

Many of the reported sex differences in the brain, studied using structural magnetic resonance imaging (MRI), are driven by intracranial volume, which has been reported to be on average 12% larger in males than in females ^12,17^. While most cortical and subcortical brain regions appear to have significantly larger volumes in males than in females and females present with greater overall cortical thickness, Ritchie et al. demonstrated that the effect of these differences decreases when the analyses are controlled by intracranial volume ^18^. A common metric used for adjustments (also referred to as normalization) is the total intracranial volume (TIV), as it is assumed to be constant across the individual’s lifespan and is not affected by aging or pathological processes. In recent years, several studies have utilized correction approaches to account for the confounding effect of brain size on sex differences, with a focus on accurate classification of females vs males based on structural MRIs beyond brain size ^19,20^. However, these results were based on prediction accuracy (i.e. classifying males versus females) rather than devising a gold standard benchmark and as such, the biological implications of disentangling the impacts of sex and brain size on neuroanatomical associations remain to be explored. Furthermore, while employing correction methods removes (some) of the TIV-related differences, each method might also introduce different types of bias when assessing the potential presence of sex differences in normalized values.

Another important question related to brain size is the scaling relationship between total and regional volumes, referred to as brain allometry. Studies on brain allometry suggest that distinct regions are scaled differently with respect to brain size ^21^ and this regional specificity is associated with the functional and biological properties of the brain ^22^. Furthermore, larger brains are not simply a scaled version of smaller ones ^23^. However, to our knowledge, there are no studies that investigate the effect of aging on brain allometry, nor on how this scaling property is different between females and males in adult lifespan.

In this work, taking advantage of the large sample size provided by the UK Biobank (UKBB) dataset, we employed a novel approach (i.e., matching) to address these questions. We investigate the presence of sex differences in samples that are matched by intracranial volume, essentially removing the impact of intracranial volume differences from the assessments. We further used the same matching approach to investigate the effect of aging on brain allometry, and whether allometric properties differ between sexes. Finally, we explored the effect of head size on perceived sex differences in cognitive performance across several cognitive tests.

## Results

We used structural neuroimaging (T1-weighted MRIs) and sociodemographic information from the UKBB dataset (see Methods). We extracted regional volumetric information based on the Desikan-Killiany Cortical Atlas ^24^ and the subcortical volumetric segmentations (Aseg atlas), as well as cortical volume, thickness, and surface area, using FreeSurfer version 7.4.1 ^25^. Following quality control, N = 35,732 participants (N = 19,281 females, 44-82 years) were included in this study. Using this sample, we created four subsamples to study the effect of sex beyond brain size on brain neuroanatomy: **(i)** an age and TIV matched sample of males and females with 11,294 participants (5,647 females), (referred to hereafter as matched sample), allowing us to disentangle the impact of TIV from sex differences,**(ii)** a non-matched sample of the same size (referred to as not matched sample), allowing us to investigate the results in a sample characteristic of the complete sample, but with the same statistical power as the matched subsets, **(iii)** a sample of the same size matched just by age (referred to as age-matched sample), allowing us to assess whether age differences between the two populations impact the findings regardless of the brain size, and **(iv)** a sample exaggerating the TIV difference between sexes by sampling from the opposite tails of the TIV distributions (referred to as extreme sample), allowing us to assess whether greater TIV differences across the groups will lead to further bias and inflated sex difference estimates. A higher effect size in this sample will confirm the major contribution of TIV, while consistent effect sizes will provide evidence for sex differences.

### Identifying sex differences beyond brain size

We modeled the aging trajectory of each region in the four subsamples using mixed effects linear models, accounting for age, sex, their interaction, and using the scanning site as a random effect ^26^. Figure 1.A presents the distribution of age and TIV across the matched and not-matched samples, showing that the matching process has yielded two perfectly matched subsamples. Figure 1.B. shows the relationship between TIV and total brain volume and age, confirming the lack of association between TIV and age. Based on these, the derived subsamples are not biased with respect to age and can be used for the downstream analyses. Figure 1.C. shows the distribution of the standardized model estimates across the 78 brain regions for each sample. The age estimations remained consistent across samples, however, the TIV-related differences between females and males changed the estimates for the effect of sex across samples. In the case of the extreme sample where the difference is further exaggerated, the estimates for the effect of sex on volumes are high for all regions which can mask the nuances of relative volumes and their changes over time. However, in the case of the matched sample, where females and males have the same head size, the estimates were centered around zero, indicating that although there were some sex differences in the trajectories, they are more subtle and, importantly, bidirectional. For instance, males had relatively lower volume estimates in the matched sample in bilateral superior parietal, bilateral caudal middle frontal, right postcentral, and left inferior parietal gyrus, and larger estimates for right lateral occipital and right fusiform gyrus, bilateral amygdala, and left putamen. Finally, the mixture of brain size and sex results in spurious interaction findings between age and sex, suggesting differential trajectories between sexes where after matching these effects disappear. For example in the not matched sample, ten regions (left amygdala, caudate, lingual gyrus, and precentral gyrus, right parahippocampal gyrus and ventral DC, and bilateral hippocampus and thalamus) showed a significant interaction of age and sex after correcting for multiple comparisons performed using the false discovery rate (FDR) controlling method, whereas in the matched sample, only 4 regions, namely the left caudate, the bilateral pallidum, and the left paracentral gyrus survived correction.

**Fig. 1:**
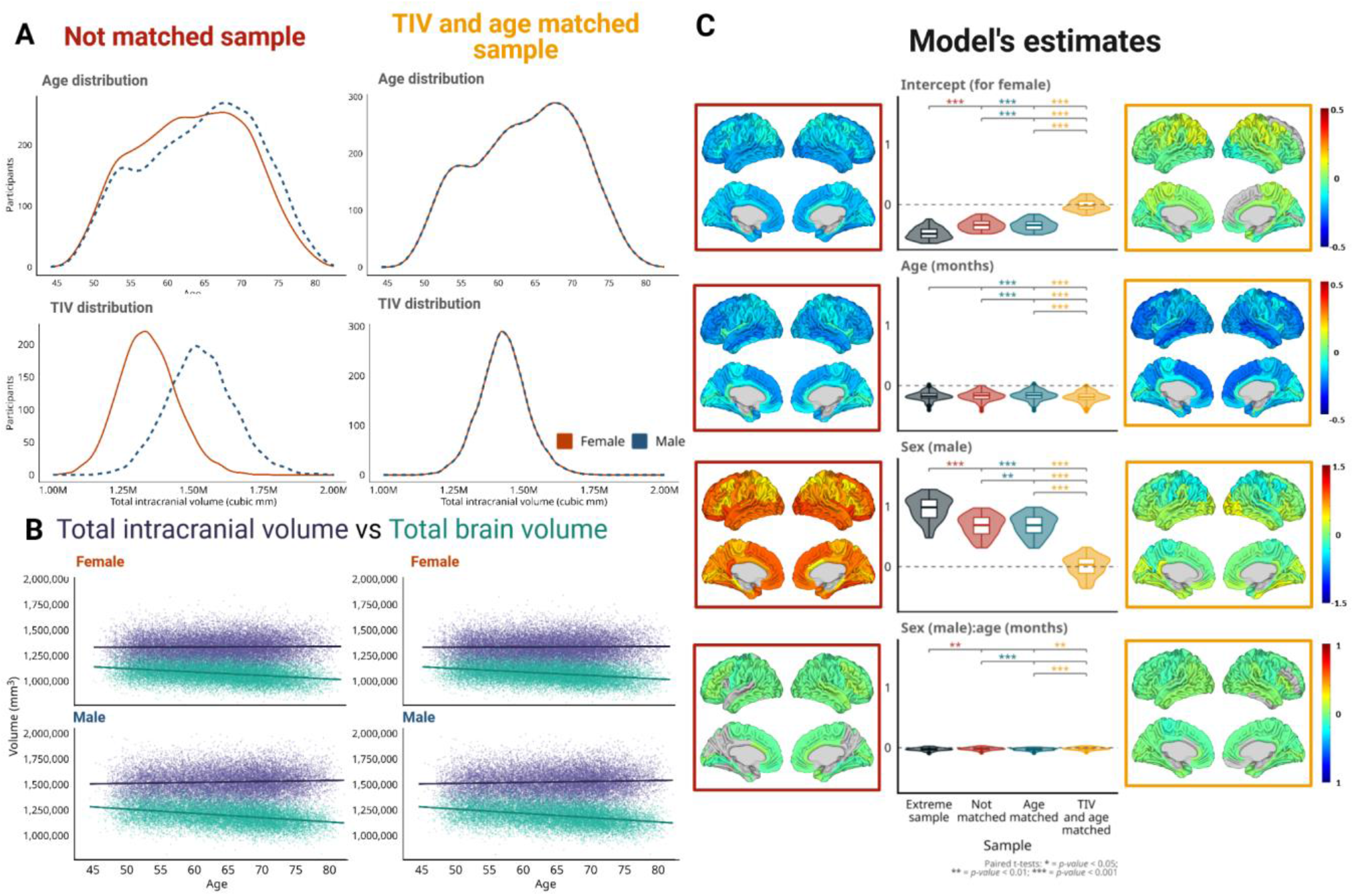
**A)** Distributions for age and TIV for not matched and matched samples. **B)** Comparison between TIV and TBV for not matched and matched samples, separated by sex. **C)** Sex differentiated aging trajectory model estimates (*Volume* ∼ *1* + *Age* + *Sex*+ *Age:Sex*). The first and third columns show the model estimates projected onto the brain for the not matched and matched samples, respectively. The middle column shows the distribution of the same estimates, including those of the subcortical regions. Note that the regional volumes and age are z-scored. Females were used as the reference group in our model and therefore, the “sex” estimation is for males compared to females.

To examine whether there might be more subtle sex-related differences that might not be captured by regional segmentations, we repeated the volumetric analyses at a vertex level. Furthermore, we performed similar vertex-wise analyses using cortical thickness and surface area measurements to disentangle the potential sources of these differences. We projected these vertex-wise estimates onto Yeo functional brain networks as well as the Von Economo cytoarchitectonic classes to examine whether the observed patterns are confined within certain cytoarchitectural and functional brain organizations or spread across the brain. Figure 2.A shows that the TIV-related differences between females and males changed the estimates for the effect of sex across samples. The pattern we observe for volumetric and surface area information is similar to that of the regional volumes, although overall the estimates are slightly larger and more variable in surface area than volume. In comparison, cortical thickness has smaller estimates and the opposite pattern, and although the estimates tend towards zero in the matched sample, females have larger estimates than males. Panel 2B shows the distribution of these estimates in functional networks^27^ and cytoarchitectonic classes^28^ in the matched sample. Interestingly, in the case of the functional connectivity networks, the three measures basically follow the same pattern, although with different magnitudes, while this is not the case for the cytoarchitectonic classes in which the volume and cortical thickness are more similar whereas surface area seems to be balanced for most of the classes. The sex estimate tends towards negative values (i.e. larger in females) for somatomotor, dorsal attention, and ventral attention networks, positive values (i.e. smaller in females) for visual and limbic networks, and minimal differences for frontoparietal and default mode networks. For the cytoarchitectonic classes, sex difference estimates for surface area are balanced for all classes, except for the secondary sensory (SS) and insular cortex, which tend towards positive values. For volume and cortical thickness, the sex estimate tends towards negative values for the primary motor and sensory cortices (OM and PS) and the association cortices (AC1 and AC2), while presenting higher values for the limbic regions and insular cortex, and having divergent patterns for the SS in which volume has more positive values while cortical thickness is balanced.

**Fig. 2:**
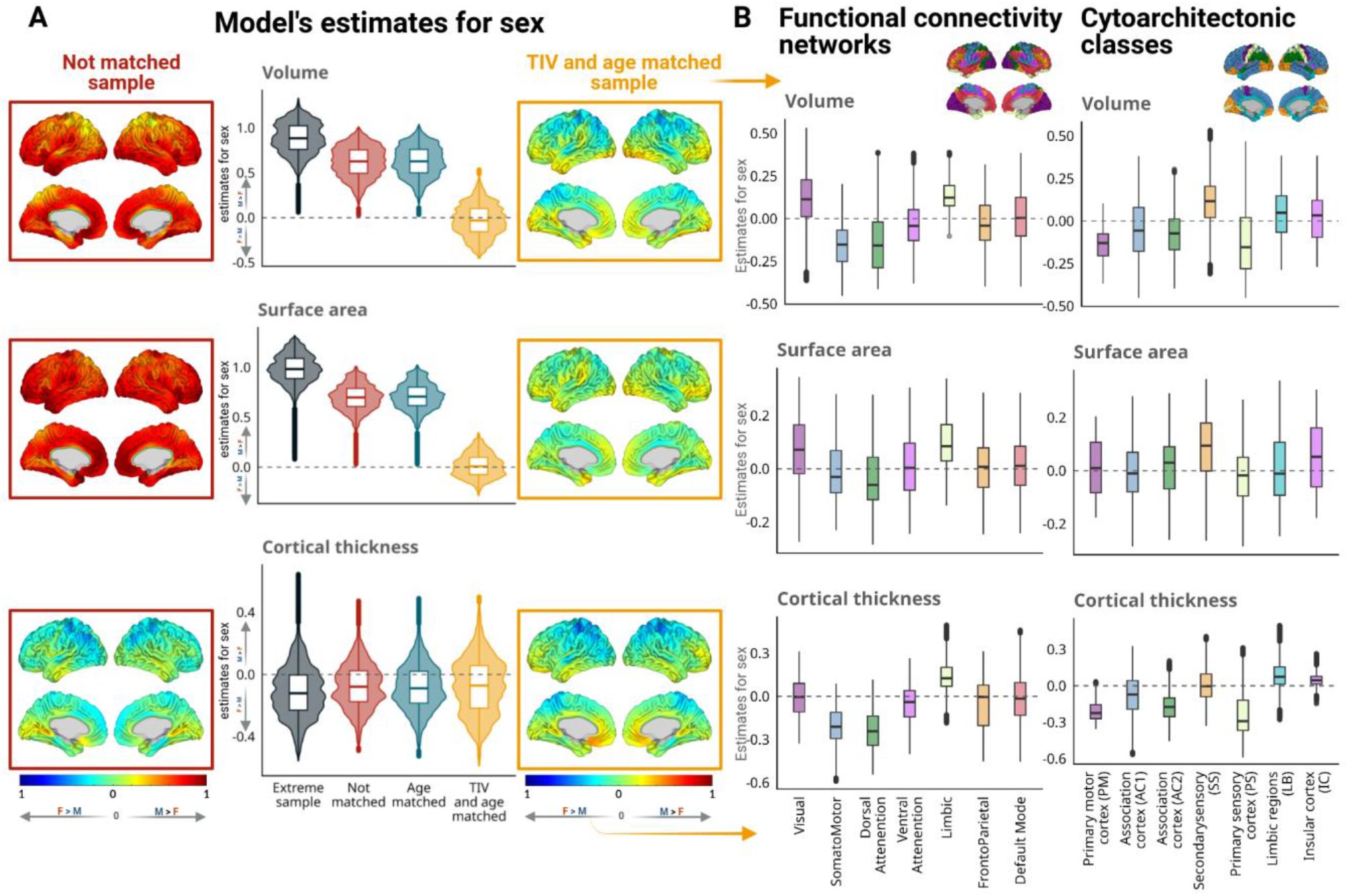
**A)** Vertex level sex estimates for the sex differentiated aging trajectory model estimates (*measure* ∼ *1* + *Age* + *Sex*+ *Age:Sex*). The first and third columns show the model estimates projected onto the brain for the not matched and matched samples, respectively. The middle column shows the distribution of the same estimates. Note that females were used as the reference group in our model and the “sex” estimation is for males compared to females and that the vertex-wise measures and age was z-scored. **B)** Sex estimates for the model projected onto functional networks and cytoarchitectural classes.

### Sex differences in the adult brain allometry

To explore the potential sex differences in the scaling relationship between regional and total brain volumes (i.e. allometry), we calculated allometry measures by modeling regional volumes as a function of total cerebral gray matter volume using a sliding windows approach and separating the samples by sex. To assess the potential impact of aging and sex differences on allometry, we then modeled the resulting allometry estimates as a function of age, sex, and their interaction. Here, the model intercept indicates the estimated allometry value for the reference group (i.e. females), and sex estimates indicate the relative differences between the estimated allometry between males and females.

The obtained allometry maps were similar in terms of their general patterns to previous literature ^22^, derived based on developing and young adult populations (i.e. ages ranging between 5-25 and 22-35 years, respectively) and regressing out the impact and sex differences. Importantly, the allometry estimates were consistent across samples, suggesting that allometry is not heavily impacted by head size variation in the adult population. Along the same lines, the magnitudes of the sex estimates were overall relatively small and consistent between our subsamples, suggesting minor sex differences in allometry, with bilateral accumbens and posterior cingulate areas as well as left caudal middle frontal regions having the largest estimated allometry differences between sexes in the matched sample (with higher values in females).

### Behavioral data

To assess the impact of head size (i.e. TIV) on the perceived sex differences in cognitive performance, we selected the results of a set of tests from the available UKBB information (for more details, see methods) and computed the effect sizes of the differences in results between males and females, both in the matched and not matched samples (Figure 4). In all cases, higher scores represent better performance (variables with the reverse relationship were multiplied by -1). Positive effect size values indicate that females had higher scores (i.e. better performance) than males. For the matched sample, the effect sizes ranged between -0.19 (the mean time to correctly identify matches in the reaction time test, which evaluates processing speed) to 0.15 (the fluid intelligence score, which assesses verbal and numerical reasoning). Meanwhile, for the not matched sample, the effect sizes ranged between -0.22 (the percentage of solved to total viewed puzzles in the tower rearranging test, which evaluates executive functioning) to 0.1 (the mean time spent answering each puzzle in the matrix pattern completion test, which assesses non-verbal reasoning). Overall, the cognitive domains that showed a change in direction in the effect size estimations while comparing the two samples were “verbal and numerical reasoning” and “working memory”, and in both cases, the specific scores that showed the greatest shifts, had the highest correlations with TIV (r = 0.17, p < 0.001 for the fluid intelligence score and r = 0.14, p < 0.001 for the maximum numbers of digits remembered correctly). Further, for the matrix completion test which evaluates non-verbal reasoning, the percent of puzzles solved followed the same pattern as the fluid intelligence test while also having a higher correlation with TIV (r = 0.13, p < 0.001), however, the mean duration spent answering each puzzle did not have that trend and had a much lower correlation with TIV (r = -0.05, p < 0.001). Taken together, these results suggest that the sex differences that are observed in non matched samples in these domains are largely driven by TIV differences between males and females.

To further examine the changes in sex difference estimates between the matched and non matched sample, we modeled the effect of sex and regional volume on fluid intelligence score. The model’s intercept was positive for all the regions in the matched sample, however it was negative for most of them in the not matched one (70 out of 78, with only the left ventral DC and superior frontal, and bilateral lateral-orbitofrontal, superior-temporal and superior-temporal gyrus, and bilateral insula). In both samples, age had a negative relationship with the fluid intelligence scores for models across all the regions. Meanwhile, the main effect of sex followed the opposite pattern, with all of the regions having a negative estimate in the matched sample (all of which were significant after FDR correction), but only the right insula being negative in the not matched sample (and not significant after FDR correction). Regarding the main effect of regional volume on fluid intelligence scores, all the estimates were positive (all corrected p-values < 0.05 in both samples), however, the estimates were larger for the not matched sample (d = 2.2). Finally, the interaction between sex and regional volume was only significant for 46 regions in the not matched sample and this effect disappeared in the matched sample for all regions.

### Additional validations

To investigate whether the matching process might have introduced potential biases in the subsets that might impact the results, we compared the socioeconomic and educational variables available from the UKBB between the participants that were included in the matched sample and those that were not. The matched sample did not have a statistically significant difference with the rest of the sample regarding educational level (*X*^2^ (11, *N* = 35,732) = 18.41, *p* = 0.07), or income (*X*^2^ (7, *N* = 35,732) = 12.213, *p* = 0.09). The educational level was assessed via a composite variable considering UKBB data fields 6138 (“Qualifications”) and 845 (“Age completed full time education”), and income via field 738 (“Average total household income before tax”). To ensure presence of subtle motion-related errors did not impact the findings, consistent analyses were also repeated using the FreeSurfer-based Euler number as a proxy for motion during scanning ^29^. There were no significant differences between these results and the models that did not account for motion (supplementary materials). Note that cases with moderate to severe motion levels were already excluded during the visual quality control step.

## Discussion

In this work, we assessed the impact of head size on the estimation of sex differences in brain regional volumes as well as cortical volume, surface area, and thickness and its relation to canonical functional networks and cytoarchitectural classes. We further examined sex differentiated aging trajectories across these measures, sex differences in allometry, and the distinct impact of sex and brain size on cognitive tests. Our results suggest the estimated sex differences in aging models substantially decrease when females and males have the same head sizes and that more subtle patterns of sex difference emerge once the impact of head size is removed. Our allometry analyses suggest that allometry is not heavily impacted by head size and that the sex differences are present but subtle. Regarding the cognitive performances, we found that head size might impact the performance in “verbal and numerical reasoning”, “working memory”, and “non verbal reasoning” domains previously perceived as sex differences.

Consistent with the literature ^3,30^, we found a decrease in cortical and subcortical gray matter volumes as participants aged. We also found sex differences in the estimated trajectories in all samples, however, as hypothesized, the differences for the exaggerated sample were greater than those of the age matched and not matched samples. The estimates for the age matched and not matches samples were in turn greater than those of the matched sample, suggesting that a significant portion of the differences reported in non-matched/adjusted samples is due to head size differences between the sexes ^17,18,20,31,32^ and not the impact of sex as a biological variable. This can also be interpreted that the main avenue for sex differences’ effect on regional brain differences is mediated through its impact on head size.

To our knowledge, our analysis is the first to explore the effects of age (across adult lifespan) and sex in brain allometry. Previous studies demonstrate that different brain regions scale distinctly with brain size and that these natural variations are related to brain functioning at distinct scales (*i*.*e*. at the micro and macroscopic levels) ^22^. Our results were similar to these findings (based on the HCP results from Reardon et al.), i.e. the intercept estimate maps showed regions with negative scaling mostly in limbic areas (and subcortical regions in our case) and positive scaling in frontal and parietal regions (Figure 3.C). The scaling relationship between regional and total volumes has been used for developing adjustment methods ^33^, and neuroanatomical norms ^21^. Our results suggest that overall, the aging process does not change the scaling relationship between regional and total brain volumes. Furthermore, this relationship is similar in males and females. The similarity of the results between non matched and matched samples also suggest the allometry follows a similar pattern across brain sizes and the estimates based on brain sizes included in our matched sample generalize to smaller and larger brains as well.

**Fig. 3:**
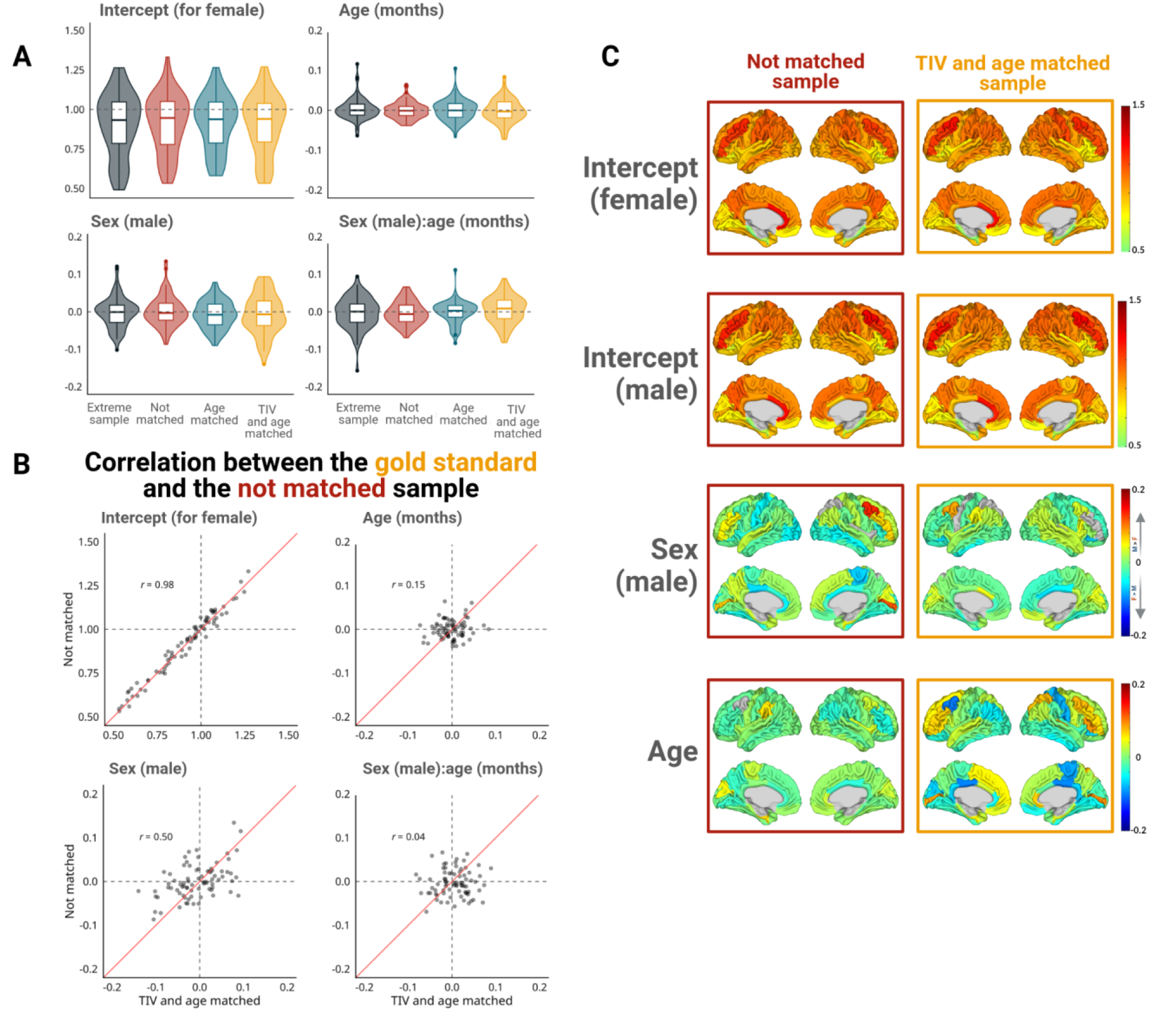
Sex differentiated aging trajectory model estimates (*allometry* ∼ *1* + *Age* + *Sex* + *Age:Sex*) for the allometry estimates for each region (*log*_*10*_*(regional volume)* ∼ *log*_*10*_*(total brain volume)*) **A)** Distribution of the estimates, including subcortical regions. **B)** Correlation between model estimates in the matched sample (gold standard) versus the not-matched sample. **C)** Model estimates projected onto the brain for the not matched and matched samples, respectively. Note that females were used as the reference group in our model and the “sex” estimation is for males bias compared to females and that age was z-scored.

**Fig. 4:**
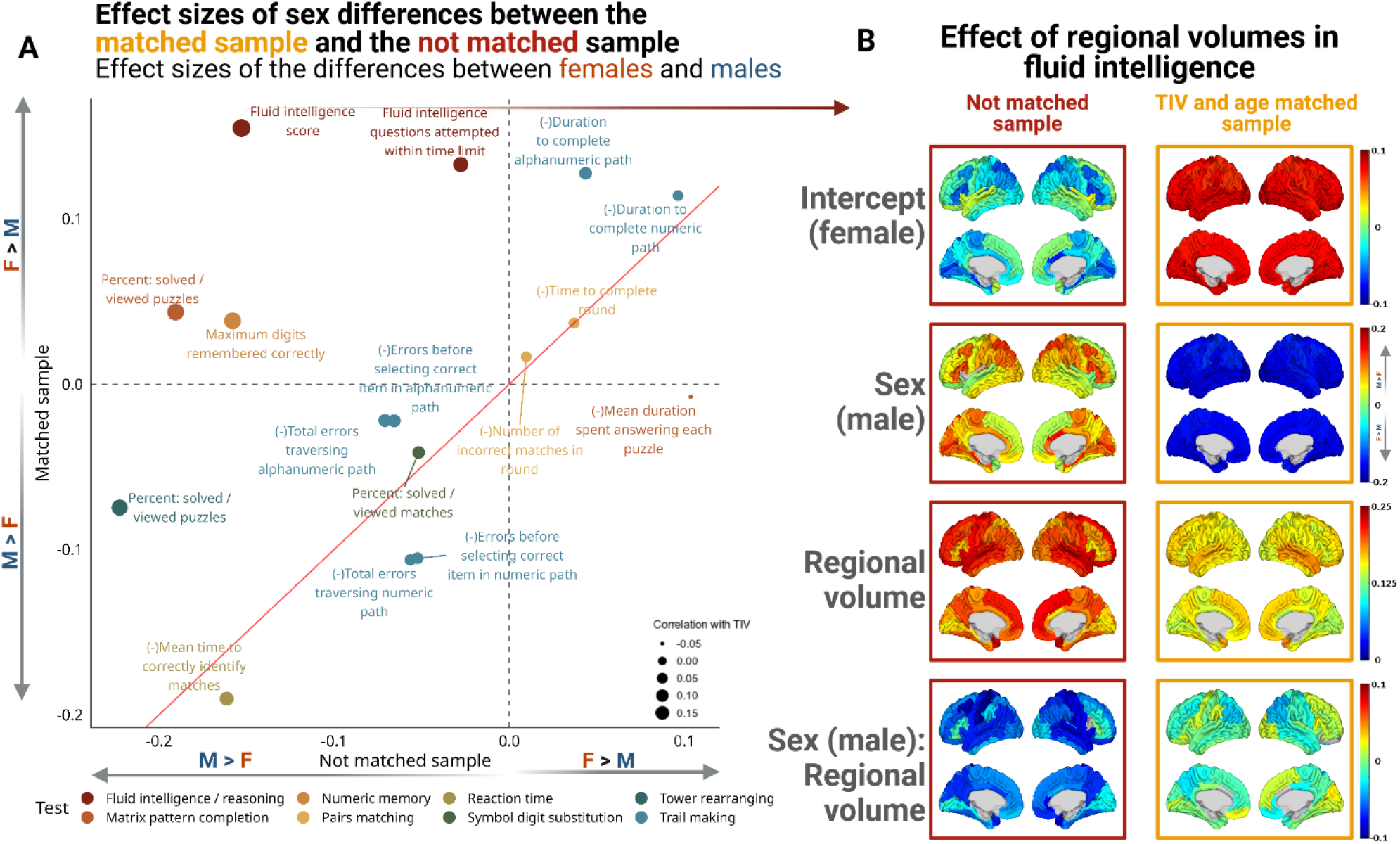
**A)** Effect sizes of the performance difference between females and males in cognitive tests in the matched and the non-matched samples. Each color represents a test and each point a score. The sizes of the points represent the correlation between the test results and TIV for all the subjects, with values ranging from -0.05 for “Mean Duration spent answering each puzzle” in the Matrix pattern completion test to 0.17 for the “Fluid intelligence score”. **B)** Model estimates assessing the effect of sex and regional volume on fluid intelligence (*Fluid intelligence* ∼ *1* + *Volume* + *Sex* + *Volume:Sex* + *age)* in not matched and matched samples. Note that females were used as the reference group in our model and “sex” estimation is for males bias compared to females and that regional volumes, fluid intelligence scores, and age were z-scored.

We explored the relationship between performance on cognitive tests, sex, and head size. In this context, sex differences from a behavioral perspective are usually studied without considerations for head size differences (or other biological variables), however, we wanted to disentangle the contributions of these two factors. Consistent with prior literature ^34^, our findings suggests that i) cognitive functioning in older age depends partially on maximal brain development (i.e. head size), ii) the contributions of maximal and current brain volumes differ among cognitive domains, and iii) that TIV is related to semantic memory, executive functioning, and spatial ability, but not to episodic memory. Congruent with what was reported by Fawns-Ritchie & Deary, females had better performance in the pairs matching test and worse in the tower arranging test. Furthermore, as previously reported, we found males had higher scores in matrix pattern completion in the not matched sample, however, this trend reversed in the matched sample, further reinforcing the notion that the observed differences are due to head size and not sex^35^. In our study, the effects of sex and regional volume in fluid intelligence scores (as the main example changing effect size between our matched and non matched sample), we found that the effects of sex were reverted between samples, that the effect of regional volume was higher in the not matched sample and that the interaction between these two factors was only significant in the not matched sample.There is a growing appreciation of the effect of omitted variable bias in establishing the causal effects of variables with highly confounding factors associated with them, as is the case with studying sex differences. Our findings further emphasizes these aspects and the importance of considering relevant biological factors in studying sex differences in behavioral domains.

Our study benefits from several strengths that enhance the reliability of our findings. We had access to a large dataset comprising more than 30,000 participants, which not only provided us with sufficient statistical power for our analyses, but also had enough range and variability to allow for perfectly matched subsamples with more than 10,000 participants. Furthermore, we relied on the results of visually quality controlled derivatives of our extensively validated in-house pipeline for the matching process, ensuring the accuracy of the TIV estimates. Other sources of error (presence of motion, incomplete field of view, incidental findings) were also flagged during the same quality control process and these cases were excluded from the analyses.

Our study is not without limitations. The UKBB dataset only includes participants older than 45 years old, and as such, our samples do not include information from younger adults and thus might not be representative of the entire life span. Furthermore, the overall UKBB sample had a 5.47% invitation to participation rate, and might not be fully representative of the general population ^18^. It has been reported that the sample is biased towards participants who are more likely to live in less socioeconomically deprived areas, less likely to be obese, report lower smoking rates and alcohol intake, and have fewer self-reported health conditions than non-participants ^36^. In addition, over 96% of the participants who had imaging data were Caucasian, limiting the generalizability of the findings to other ethnic groups. Future investigations in more diverse populations are therefore warranted to examine the generalizability of the observed findings. Although our matching approach provided us with a gold standard to evaluate sex differences and compare other methods against, it is worth noting that it led to restriction of the range of head sizes, diminishing the variability in the samples. Finally, although the cognitive tests used by the UKBB correlate with well-validated standard tests (moderate to high correlations), they have been reported to show lower test-retest reliability than some of the tests that assess the same cognitive domains and might not capture the full range of variability in the cognitive functioning^35^.

## Methods

### Datasets

#### UKBB dataset

Participants were included from the UK Biobank, an open-access large prospective study with phenotypic, genotypic, and neuroimaging data from 500,000 individuals recruited between 2006 and 2010 at 40–69 years old in Great Britain ^37^. All participants provided informed consent (“Resources tab” at https://biobank.ctsu.ox.ac.uk/crystal/field.cgi?id=200). The UK Biobank received ethical approval from the Research Ethics Committee (reference 11/NW/0382) and the present study was conducted based on application 46 007. Sex was assigned according to the “Sex” variable (data field 31).

### Image processing

We utilized data from the first imaging visit (instance 2). We processed the raw T1-weighted images using FreeSurfer v.7.4.1.^25^ and extracted the volumetric information. We used the Desikan-Killiany parcellation ^24^ and the Aseg atlas segmentations for the cortical and subcortical regions, respectively. Since it has been reported that the estimated intracranial volume (eTIV) from Freesurfer is biased and might not be optimal for TIV normalization ^38^, we used a widely used and extensively validated in-house pipeline based on the open source MNI MINC toolkit (https://github.com/BIC-MNI/minc-toolkit-v2) to obtain accurate TIV estimations ^39–45^. The images were preprocessed by denoising using optimized non-local means filtering ^46^, corrected for intensity inhomogeneity ^47^, and intensity normalized using linear histogram matching. The resulting images were linearly registered to the MNI152-2009c template (nine-parameters) via a hierarchical linear registration method ^39^. From the linear registrations, we obtained a scaling factor that allowed us to estimate TIV, similar to the process used by FreeSurfer to estimate eTIV. All registrations were quality controlled by two experienced raters (Y.Z. and M.D.), to ensure the accuracy of the obtained TIV estimates.

### Matching process

The final sample included data from 35,732 participants (54% females). The exclusion criteria included: unavailability of TIV, brain segmentation metrics, volumetric measures, or demographics information (e.g., age, sex, etc.), and failed visual QC of image processing and registration steps. With these, we created three subsamples including equal numbers of male and female participants for the analyses. The first subset was matched by age, calculated as months between the participant’s month (data field 52) and year of birth (data field 34), and the date of visiting the assessment center (data field 53) for the imaging visit (instance 2), and by TIV. To achieve this, we split the sample by sex. Since our original sample had more females than males (19,281 F vs 16,451 M), we randomized the samples and used an iterative process of finding a female participant whose age (in months) and TIV estimates were within a 0.02% range from the corresponding values of each male participant. Following this process, we obtained a sample of 11,294 matched participants (5,647 per sex). Similarly, we created a second subsample with the same size (i.e. 11,294) matched only by age. We created a third sample of the same size that mimicked the age and TIV distribution of the original full sample, to allow for comparisons of the results across matching strategies with the same statistical power. Finally, we created a sample that exaggerated the TIV difference between females and males by excluding all the participants that were included in the matched sample and keeping 6,647 participants per sex whose age distribution was similar to the original sample. Table 1 presents the information for all generated samples.

**Table 1:** sample description.

| Sex | N | Age range | Age (mean, sd) | TIV (mean, sd) |
| --- | --- | --- | --- | --- |
| **Full dataset** |  |  |  |  |
| Female | 19,281 (54%) | 45 - 81 years | 62.8, (7.36) | 1,336,544.0, (103,196.89) |
| Male | 16,451 (46%) | 44 - 82 years | 64.0, (7.64) | 1,525,386.9, (119,264.92) |
| **TIV and age matched sample** |  |  |  |  |
| Female | 5,647 (50%) | 47 - 80 years | 63.3, (7.06) | 1,426,776.1, (83,886.29) |
| Male | 5,647 (50%) | 47 - 80 years | 63.3, (7.06) | 1,426,823.3, (83,880.34) |

| Age matched sample |  |  |  |  |
| --- | --- | --- | --- | --- |
| Female | 5,647 (50%) | 45 - 81 years | 64.0, (7.67) | 1,336,051.7, (103,836.41) |
| Male | 5,647 (50%) | 45 - 81 years | 64.0, (7.67) | 1,525,577.3, (117,849.54) |
| Not matched sample |  |  |  |  |
| Female | 5,647 (50%) | 46 - 81 years | 62.8, (7.39) | 1,338,532.7, (103,282.83) |
| Male | 5,647 (50%) | 44 - 81 years | 64.0, (7.67) | 1,524,394.1, (118,165.63) |
| Extreme sample |  |  |  |  |
| Female | 5,647 (50%) | 45 - 81 years | 63.6, (7.02) | 1,297,634.6, (85,362.83) |
| Male | 5,647 (50%) | 44 - 82 years | 64.4, (7.95) | 1,574,611.6, (101,046.49) |

### Modeling

The following linear mixed effect models were consistently performed on the regional volumetric information:

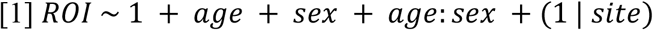

Where *ROI* denotes the region of interest, *age* is the age at acquisition time in years (data field 21003), *sex* is the participants’ sex (data field 31), and *site* represents the assessment center where image collection was conducted (field 54). We then examined the effect sizes for each model and used paired t-tests to assess the differences across subsamples. All the statistical analyses were performed using the R statistical language.

Similar analyses were completed at a vertex level, using vertex-wise FreeSurfer volume, thickness, and surface area measures. To assess whether the observed patterns of sex differences were differentiated across known brain networks, we extracted the vertex-wise volume, cortical thickness, and surface area measures across Yeo 7 functional networks^27^ as well as the Von Economo cytoarchitectural^28^ classes and compared the regional estimates across different classes. Vertex-wise analyses were performed in MATLAB R2023a.

### Allometry

To evaluate whether the allometric scaling of different brain regions are sex and age differentiated, we calculated an allometry value for each region, for each sex, in each sample. To evaluate the effect of aging, we used a sliding window approach considering the participants’ age in months at the time of scanning. Each window consisted of 5 years so for each analysis. We considered 2.5 years before and after each continuous month to split the sample. We extracted the allometry values using the following equation:

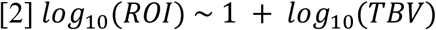

Where *ROI* represents the region and *TBV* is the total brain size. For TBV, we calculated the total cerebral gray matter volume by summing the cortical and subcortical gray matter values from the FreeSurfer Aseg atlas segmentation estimates. In this case, the allometry value is the estimate of the *log*_10_(*TBV*) coefficient in the regression model. Then, for each sample, we combined the values for females and males, assigning the central month within the window as the age value, and performed the following linear models:

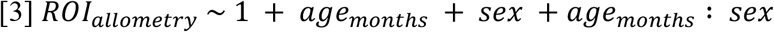

### Behavioral information

We extracted the cognitive performance information for the same instance as the imaging data from the UKBB (category 100026, see https://biobank.ndph.ox.ac.uk/showcase/label.cgi?id=100026 for additional details). Data cleaning process included selecting the variables that had numeric data as outcomes, averaging the information of different runs or “steps” in the case of repeated experiments, and negating some variables to match the direction of all variables (i.e. higher values corresponding to better performance). We used information from the data fields presented in table 2.

**Table 2:**
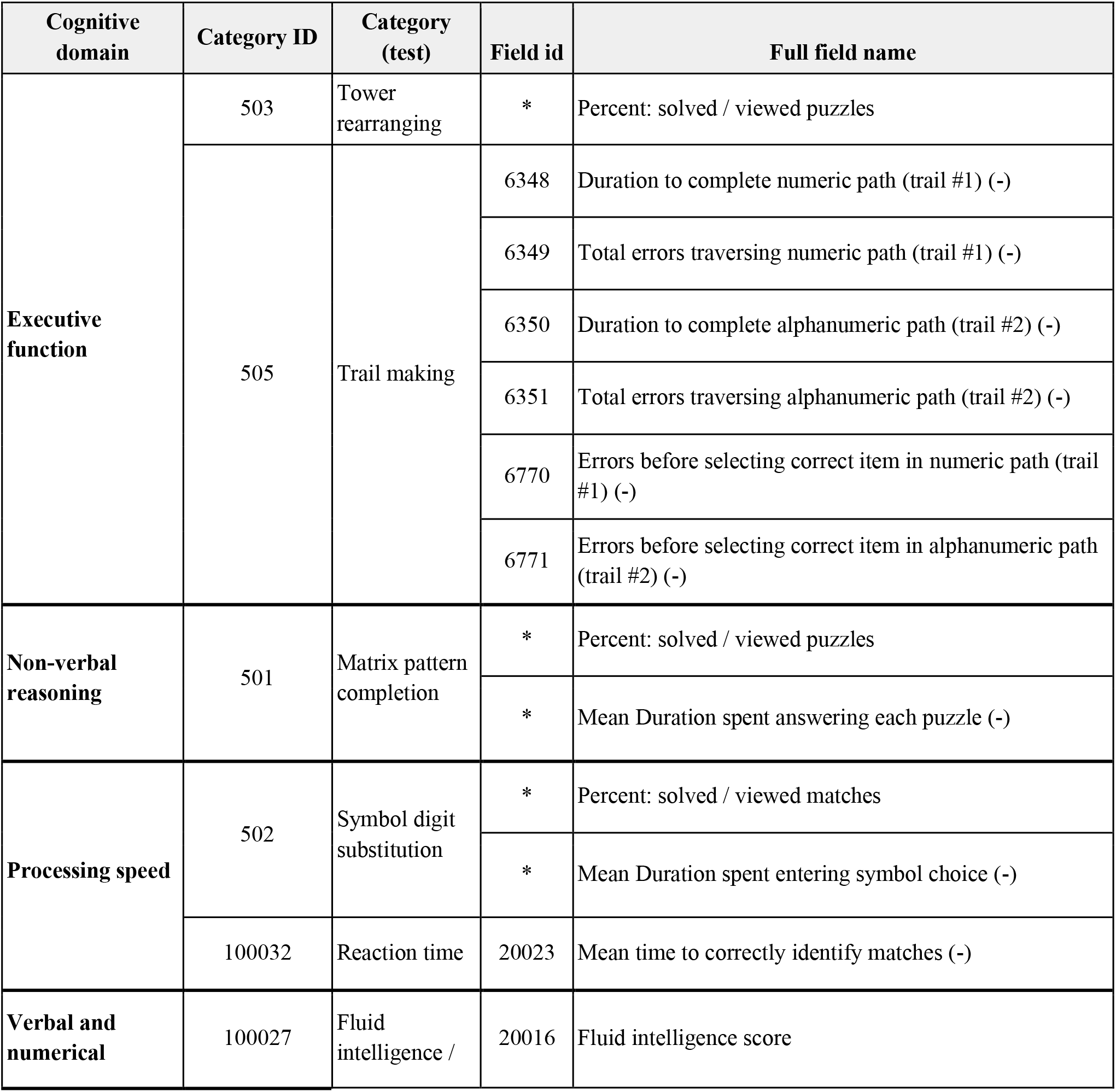
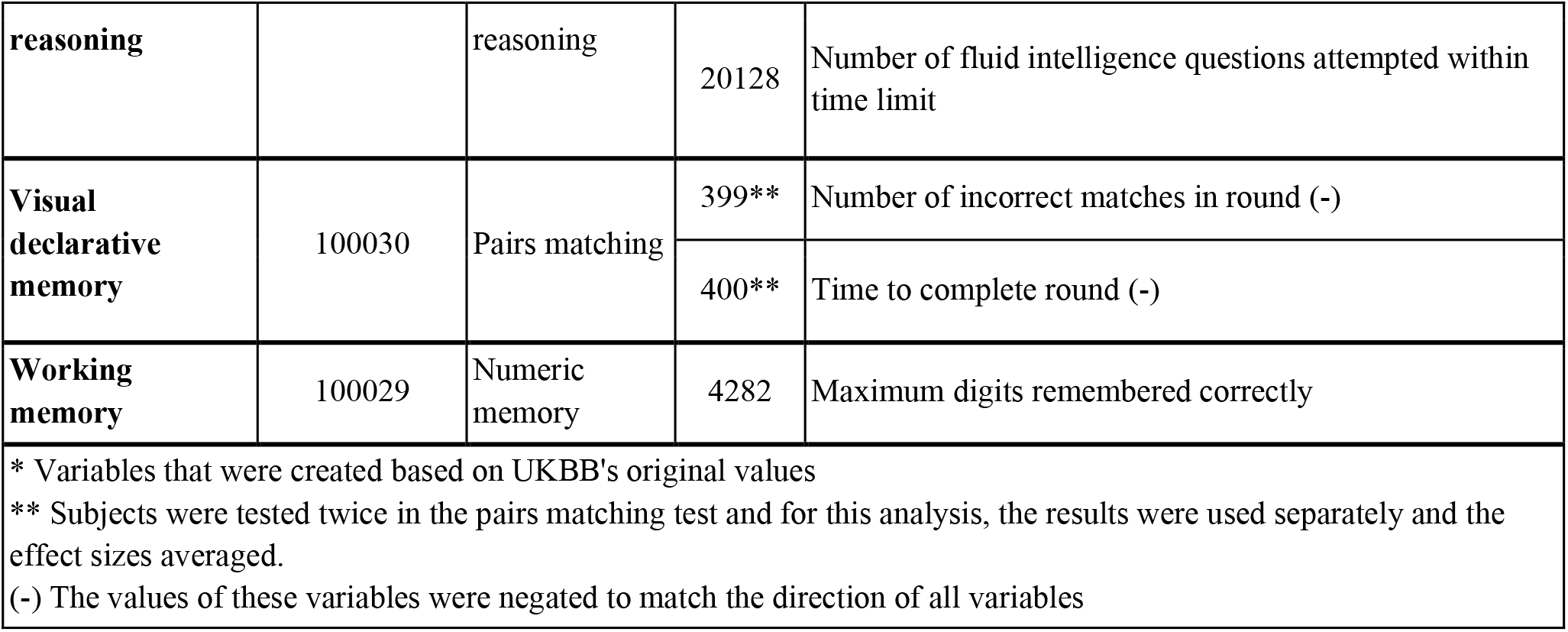
list of cognitive tests included.

For each variable and sample, we calculated and compared the effect size of the difference in the results obtained by females versus males. It is worth noting that not all participants completed all cognitive tests. The smallest percentage of participants that completed the cognitive tests for each sex was 63% (see supplementary materials for more details).

We further evaluated the relationship between regional volumes, sex, and fluid intelligence scores using the following linear regression model:

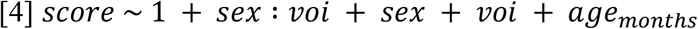

To assess whether the results were influenced by potential differences in education levels, we repeated the same analyses, adding education as a covariate, and obtained similar results.

## Acknowledgments

This research was conducted using the UKBB Resource under approved application 46 007. We thank the UKBB participants and team for their work in collecting, processing, and disseminating these data for analysis. The authors also acknowledge use of Compute Canada (https://alliancecan.ca/en) resources for performing the image processing analyses in the presented work.

